# Vessel Metrics: A python based software tool for automated analysis of vascular structure in confocal imaging

**DOI:** 10.1101/2022.12.22.521670

**Authors:** Sean D. McGarry, Cynthia Adjekukor, Suchit Ahuja, Jasper Greysson-Wong, Idy Vien, Kristina D. Rinker, Sarah.J. Childs

## Abstract

Images contain a wealth of information that is often under analyzed in biological studies. Developmental models of vascular disease are a powerful way to quantify developmentally regulated vessel phenotypes to identify the roots of the disease process. We present vessel Metrics, a software tool specifically designed to analyze developmental vascular microscopy images that will expedite the analysis of vascular images and provide consistency between research groups.

We developed a segmentation algorithm that robustly quantifies different image types, developmental stages, organisms, and disease models at a similar accuracy level to a human observer. We validate the algorithm on confocal, lightsheet, and two photon microscopy data in zebrafish. The tool accurately segments data taken by multiple scientists on varying microscopes. We validate vascular parameters such as vessel density, network length, and diameter, across developmental stages, genetic mutations, and drug treatments, and show a favorable comparison to other freely available software tools. Vessel Metrics reduces the time to analyze experimental results, improves repeatability within and between institutions, and expands the percentage of a given vascular network analyzable in experiments.

**Summary statement:** Vessel Metrics is an automated software tool designed to standardize and streamline the analysis of vascular microscopy images.

## Introduction

Quantitative analysis of vascular imaging in animal disease models can give insight into the mechanisms of human vascular disease. Vascular images contain an enormous amount of information that is often underutilized due to the time needed to manually analyze the data and the lack of familiarity with software tools to automate the analysis. Although large vessels are easily quantified, microvasculature continues to be both difficult to image and analyze. Accurate analysis of vessels is critical for characterizing changes in vascular structure as a result of disease and can provide critical information on disease state and progression, or the efficacy of a potential treatment. Inter-user variability of manual measurements can sometimes be larger than the effect size being studied due to low sampling. Furthermore, image parameters as well as protocols for taking quantitative measurements can vary between institutions, even if the same model organism and disease are being studied. There is a need for software to standardize the analysis of complex vascular imaging data to ensure repeatability and reproducibility between institutions.^1^

Automated analysis of vascular images is most often developed for human retinal images due to the existence of manually labeled gold standard datasets and direct relevance to patient treatment.^2^ Retinal images show a solid, homogeneous contrast within the vessels, low raw vessel signal, and retinal tissue with signal nearly as intense as the vessels. Reproducibility is also critical in fields beyond human retinal imaging, but current software tools and models designed to work on human retinal data will not work on confocal microscopy data which typically have irregular vessel signal localized to the vessel wall, but not the interior of the vessel, and cell-to-cell differences in transgene expression or antibody staining. Microscopy images are typically black and white and can also have an unwanted fluorescent background. As a result of differing image collection and methodology, accurate segmentation tunable for different imaging modalities and acquisition parameters is critical for a general use image processing pipeline.

The process of analyzing vascular imaging first involves segmenting the image data, vessels will be labeled with one value and non-vessels with another to give a binary mask of the structure of the vascular network. The primary purpose of generating this segmentation mask is to enable automatic analysis of structure with no user variance introduced. This segmentation can happen in multiple ways, manipulation of color and shape properties or with a machine learning-based approach which are the most common.^3^ Deep learning methods have been favored due to higher performance on a dataset by dataset basis^4–6^;however, morphologically based algorithms have advantages in that they follow a concrete set of rules that are easier to intuitively understand and adjust, resulting in a more generalizable tool. Additionally, morphological algorithms require less data to train compared to both machine learning and deep learning algorithms, which is better suited for the small sample numbers often seen in microscopy experiments.

Developmental models such as the zebrafish are commonly used as a model for vascular development and disease, while the mouse is often used for adult brain vessel imaging.^7,8^Zebrafish develop rapidly and are optically transparent, allowing for non-invasive in vivo imaging using fluorescently tagged vessel walls. Additionally, the zebrafish vasculature mirrors the human system anatomically and genetically.^9^ Despite the prevalence of zebrafish as a model, analysis of zebrafish imaging is still largely performed by hand using tools like ImageJ.^10^ Which vessels, how many measurements per image, and the methods used to extract quantitative metrics can vary between trials by the same researcher, and procedural differences between studies done in different locations can potentially create variability larger than the effect size being measured. Vessel analysis software can substantially reduce time spent manually analyzing images, increase the sample of vessels analyzed, and ensure reproducibility.^11^ Being able to sample multiple timepoints non-invasively using transgenic zebrafish and confocal microscopy is a huge advantage of the system, but it must be noted that the dynamic addition of newly forming vessels in embryonic animal models can add additional confounding signals that are difficult to segment due to them being incomplete and unlumenized even though they are marked by transgene expression.

This manuscript presents the development and validation of a new vessel analysis tool, Vessel Metrics. Vessel Metrics is a python library made to automate vascular analysis. Developed on fluorescent confocal microscopy images, Vessel Metrics aims to fill a niche in automating the analysis of vascular images in microscopy experiments across image types and animal models. We demonstrate the tools’ utility in segmenting and analyzing wild type and mutant zebrafish images, along with feasibility in murine data. Our software package can output a standardized set of vessel structure metrics indicative of disease progression. In addition, we present a fully automated vessel diameter measuring function capable of measuring every vessel in an image, which to our knowledge is not available in any of the prior commercial or open source software tools. We believe Vessel Metrics has a place in the field of vascular analysis as a multi-functional, general-purpose vascular analysis tool with a focus on confocal imaging.

## Results

### Vessel Metrics algorithm

Accurate segmentation of blood vessels is one of the most important steps in pattern analysis. Segmentation describes making a label image where pixels associated with vessels have a value of 1 and the background has a value of 0, although the underlying raw data is often highly variable. The brain vasculature of zebrafish is complex but has identifiable vessels that can be used to standardize data collection from image to image. The vessels are imaged from live embryos expressing either GFP or mCherry under vascular endothelial promoters in transgenic lines *Tg(kdrl:mCherry)*^*ci5*^ or *Tg(kdrl:EGFP)*^*la116*^. In our imaging, typical vessels in the dorsal zebrafish brain at 75 hours post fertilization (hpf) are between 2.6 and 13 μm in diameter, of which vessels less than 5 μm are typically pericyte-covered capillaries and those greater than 10 μm are smooth-muscle covered arterioles.^12^ Manipulation of signaling pathways using small molecules can dramatically alter the pattern of brain vessels, providing a test dataset for our segmentation. We collected confocal microscopy images in a z-stack covering the vessels in the entire dorsal mid- and hindbrain of a zebrafish embryos, and preprocessed images by generating a maximum intensity projection (Figure 1). Preprocessing involved standard steps in image analysis. Previously published approaches have used a rolling ball filter for background suppression^4^; however, we found a top hat filter produced nearly identical results while also reducing processing time per image from 2 minutes to 15 seconds. We then applied a smoothing filter which evens out differences in transgene expression. This was followed by a linear contrast stretch to normalize intensity differences caused by acquisition or variability between animals.

**Figure 1.**
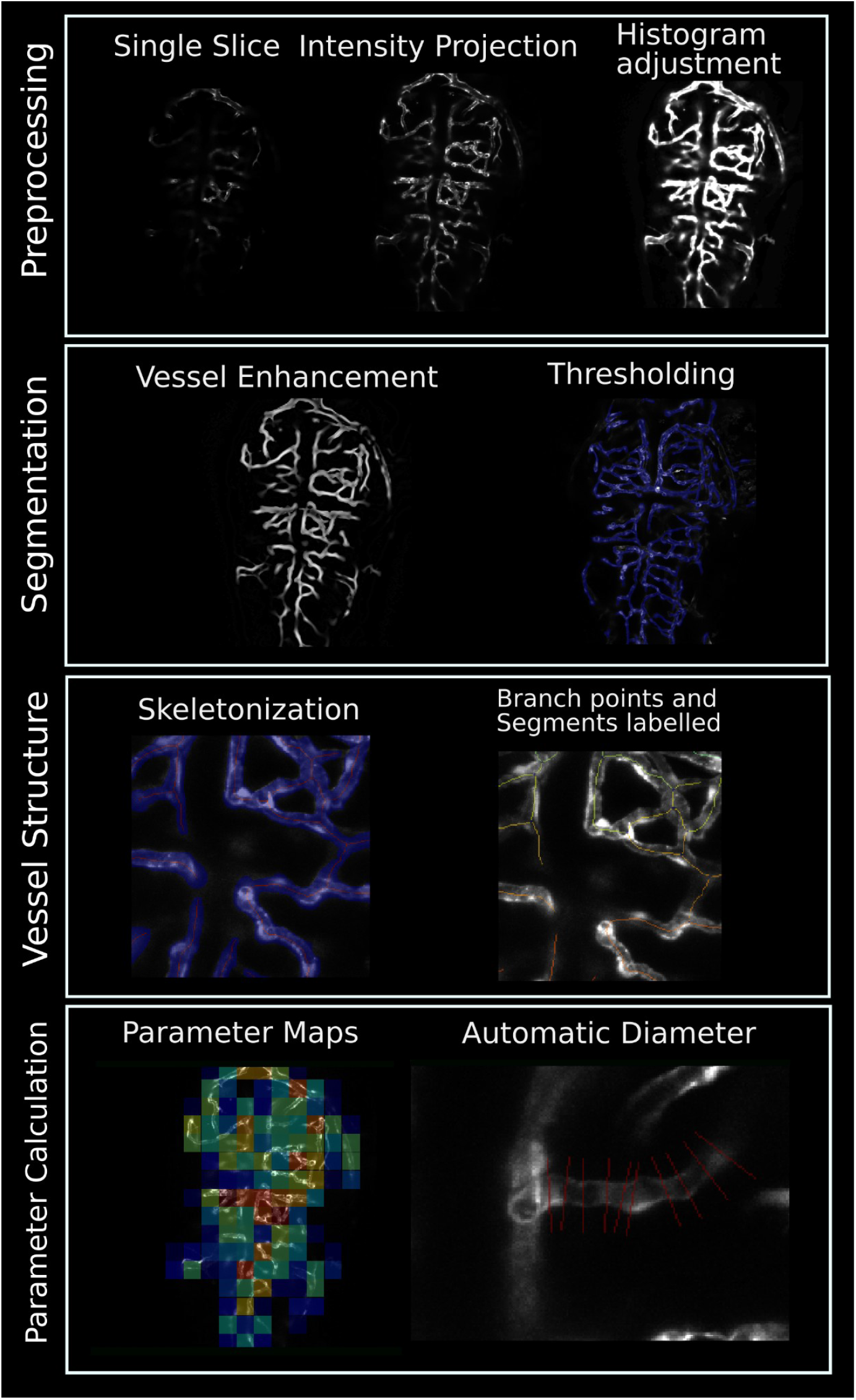
Workflow for segmentation of confocal vessel images. Preprocessing involves curating single slices into a maximum intensity projection, followed by further processing to homogenize vessel contrast and increase brightness. Segmentation involves application of one of four vessel enhancement filters (Meijering filter shown). A threshold is applied to the enhanced image to form the vessel mask. Vessel structure is determined after skeletonization of the segmented images. Extensive editing at this step removes errors. Branch points and individual segments are labeled.

**Figure 2.**
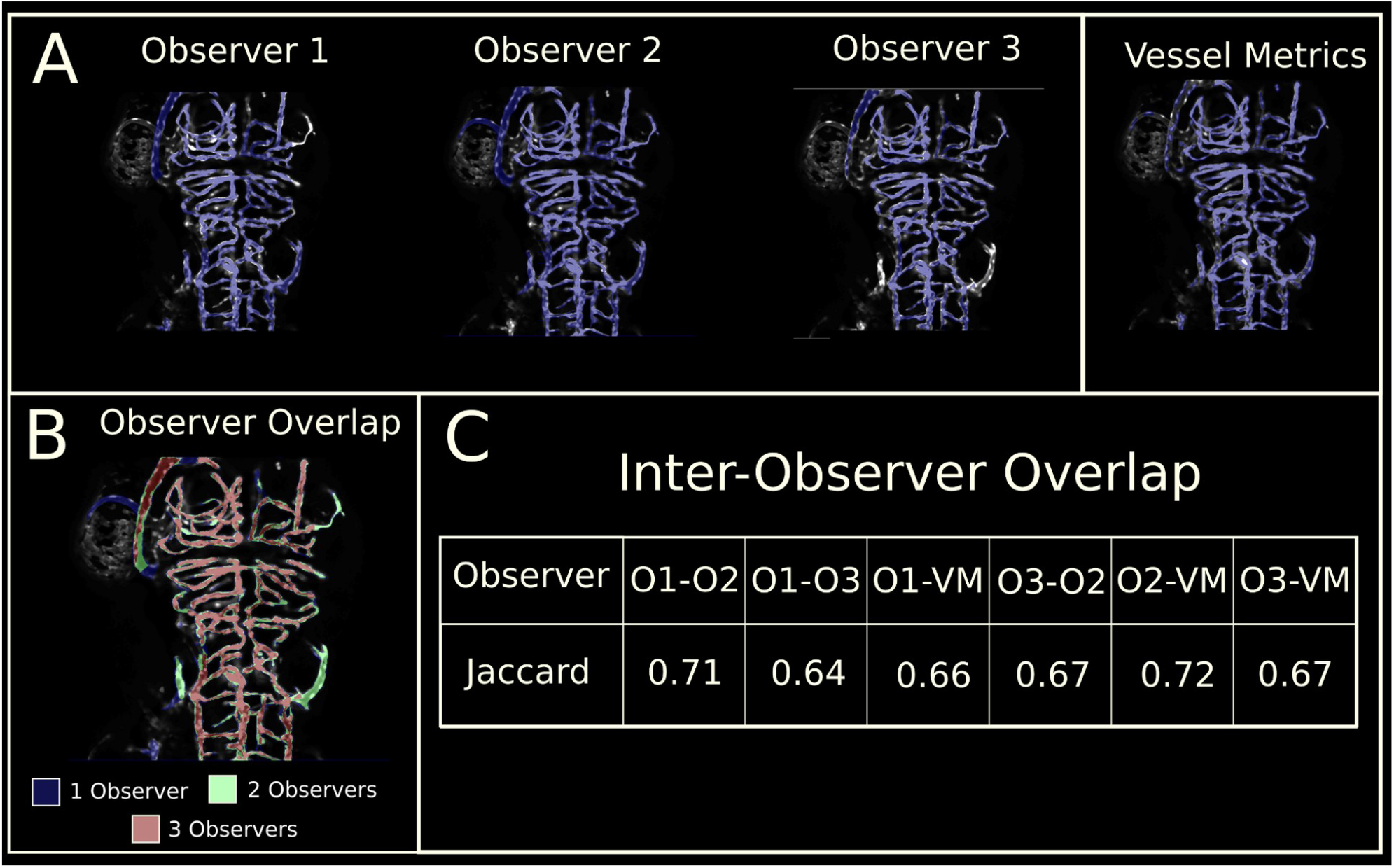
Comparison of manual user annotation and Vessel Metrics segmentation. A: Individual annotations of the 75 hpf brain vascular pattern (n = 5) from 3 observers as compared to Vessel Metrics segmentation B: Color coded agreement between observers from panel A. Red indicates all observers agree that a pixel is part of the vascular structure. C: Inter-observer overlap (n = 5). O1-O3 are observers 1-3 and VM is Vessel Metrics.

Vessel enhancement filters are employed to further brighten and homogenize vascular contrast while diminishing contrast from external tissue. Four vessel enhancement filters were tested, Frangi, Sato, Jerman, and Meijering. All four vessel enhancement filters are based on a ratio of Hessian eigenvalues. The Sato filter is mathematically the simplest with Meijering, Frangi, and Jerman adding additional terms to improve results. To compare the accuracy of different pre-processing steps, accuracy metrics were determined using the Jaccard index and a Quality factor. Accuracy metrics combine 3 vessel specific segmentation metrics: connectivity (blood vessels should form a continuous network), area (a measure of how much of the blood vessel was measured using the automated analysis vs. manual measurements) and length (the end to end distance of all vessel segments).^13^ The vessel enhancement filter with the highest Jaccard index for the test data was the Meijering filter. Using this filter with a threshold value of 40 showed a Jaccard index = 0.66 and Q = 0.7 (connectivity = 1, area = 0.76, and length = 0.92; Figure 1, Supplemental Figure 1, Supplemental Figure 2), and was used for all subsequent analysis.

### Vessel Metrics shows similar accuracy to independent manual annotations

To demonstrate how effective the automated segmentation is, we compared it to annotation by three human observers. Confocal images of 75 hpf control (DMSO) fish expressing the *kdrl:mCherry* transgene at 3 days post fertilization (dpf, n = 5) collected with identical parameters were used in this analysis. Observers manually annotated the vessel pattern from the confocal image. Inter-rater reliability was calculated as the pairwise Jaccard index between observers and the Vessel Metrics algorithm. The mean Jaccard index between pairs of human observers was 0.67, the mean Jaccard index of any human observer compared with Vessel Metrics is 0.68 suggesting that automated segmentation is as likely to agree with a human observer as another human observer.

### Automated skeletonization shows similar accuracy to manual annotation

We next optimized automated segmentation of confocal microscopy data using transgenic lines, as it poses some challenges not seen in other types of data. For instance, transgene expression can be variable between the nucleus and cell membrane, the saturation of images from microscopy data varies between users and days, and there may be a large number of angiogenic sprouts that are not lumenized in early development. In order to accurately calculate vessel parameters, vessel centerlines must first be derived using a skeletonization technique. Skeletonization produces a single pixel centerline from a binary object (our segmentation). We implemented the skimage method, which iteratively removes pixels from the boundaries of a binary object until further removal would break connectivity.

We found that without post processing, the skeleton made from our segmentation led to an overestimation of the number of segments and branchpoints. Regions containing bulbous nuclei or tortuous vessels cause burrs to appear in the skeleton, resulting in an inflated number of branch points and segments. These errors need to be differentiated from true angiogenic sprouts. We developed additional post processing steps in Vessel Metrics to correct these errors in vessel skeletonization in confocal images (Figure 3). We identified small terminal segments in the data by identifying segments thresholding with a short length and using the criteria that the short length segments were also associated only with only one branchpoint. We then eliminated these as these erroneous vessel segments. The pair of endpoints between two vessels associated with a branch point where a segment was removed is then identified. The branch point is deleted and replaced with a straight line connecting the two segment endpoints such that running the branch point detection algorithm in the future will no longer find a branch point at that location.

**Figure 3.**
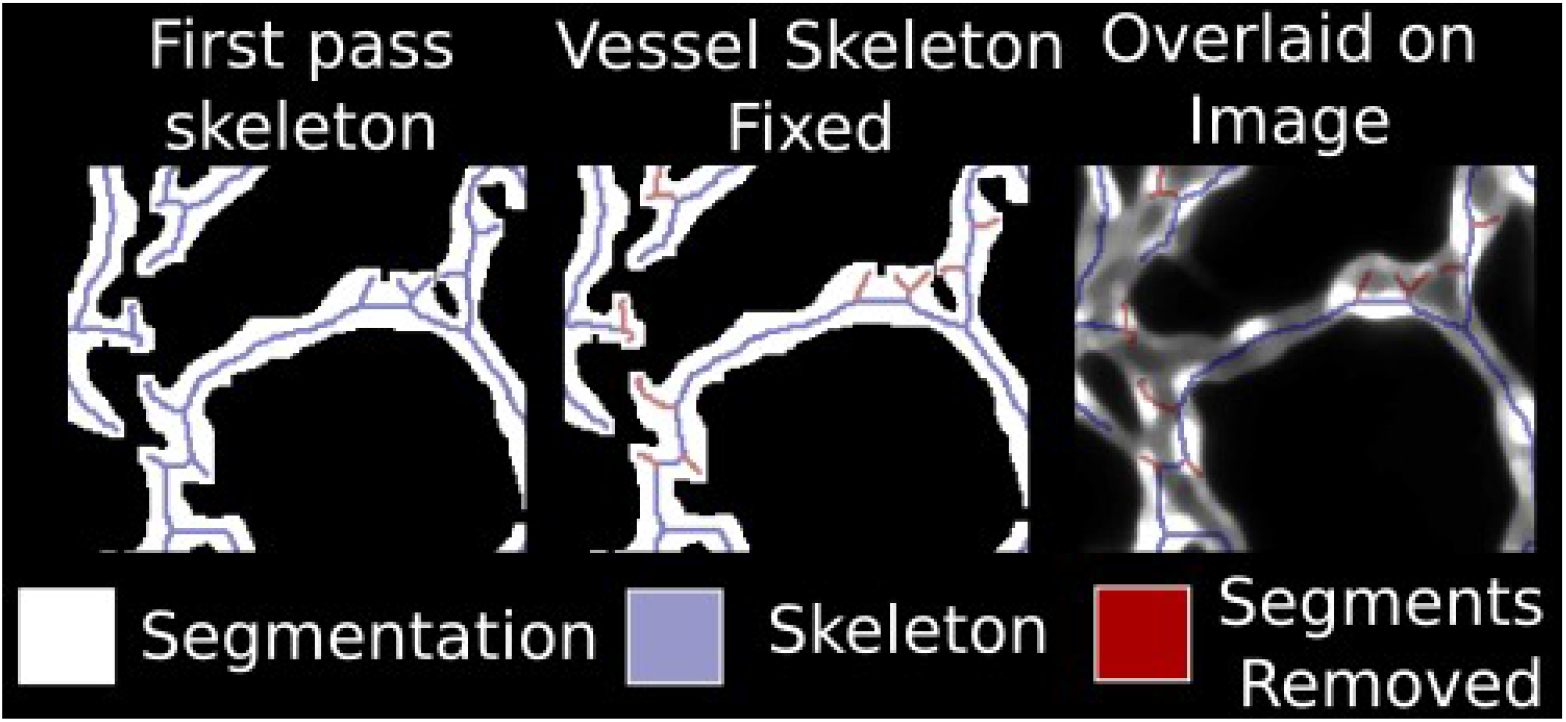
Post processing to fix errors in vessel skeletonization. Short terminal segments are pruned from the skeleton and parent branches with similar slope are connected to give an accurate representation of segment number and branch point density. Left: Raw image Middle: Vessel skeleton without post processing Right: vessel skeleton with segments removed in post processing shown in light blue.

Additionally, the raw skeletonization also over-predicted segment number as any large parent vessel with multiple child branches were considered individual segments separated by a branch point. In order to connect the segments in a parent vessel, the segment associated with each branch point is identified. The average slope of the segment is calculated using the segment endpoints. Segments associated with the same branch point with similar slopes are combined into longer single vessel segments. A similar branch point editing technique is used, a new segment is drawn between the end points and is subtracted from the old branch point, resulting in the retention of the branch point for child segments without breaking the segment.

A number of comparisons were made to test the accuracy of the segment and branchpoint count parameters. Manual measurements were made on 4 regions of interest. These manual measurements were compared to Vessel Metrics (segmentation and post processing), Vessel Metrics without post processing (to determine how effective the post processing is), and Vessel Metrics with post processing applied to a manually completed segmentation as a measure of consistency. Without Vessel Metrics post processing, we found a mean percent difference of 102% and 51% for segments and branch points respectively, indicating that without additional post processing the first pass measurements are inaccurate. After adding post processing, segment count overestimated by 26% and branch points by 15%. Repeating the same experiment with manual annotations and Vessel Metrics post processing resulted in a mean percent difference of 24% and 14% respectively, indicating consistency in the results independent of segmentation.

### Segmentation of wildtype embryos and embryos with angiogenic defects

We first tested the accuracy of Vessel Metricsby testing its ability to segment vessels in embryos with perturbed angiogenesis as compared to wildtype 75 hpf brain vessels that the program was optimized for. We exposed embryos to inhibitors of the Wnt signaling pathway (10 µM IWR-1) or VEGF pathway (10 µM DMH4) during embryonic development from 24-75 hpf, a window previously shown to inhibit brain vessel angiogenesis^14–16^. Alternatively, to detect an increase in vessels, animals at 3 dpf were compared to animals at 5 dpf where extensive angiogenesis has taken place.

Vessel Metrics default settings were applied to two embryos in each of three conditions and controls (vehicle DMSO-treated), 5 dpf vs 3 dpf, IWR-treated or DMH4-treated as compared to the control data taken at 3dpf (Table 2). We find the Jaccard, Q value, connectivity area and length are indistinguishable in the wildtype or non-standard data, suggesting that the segmentation is equally effective across manipulation and stage. Thus, Vessel Metrics will function on a broader range of data than the specific type of data it was created on.

**Table 1.** Manual parameter validation for branch point number and segment number. Manual is the manually counted number of segments or branch points. Vessel Metrics is the final

| Segment number |  |  |  |  | Branch point number |  |  |  |
| --- | --- | --- | --- | --- | --- | --- | --- | --- |
| Sample | Manual | Vessel Metrics | Vessel Metrics, hand drawn label | Vessel Metrics, no post processing | Manual | Vessel Metrics | Vessel Metrics, hand drawn label | Vessel Metrics, no post processing |
| 1 | 40 | 50 | 50 | 74 | 25 | 26 | 26 | 38 |
| 2 | 35 | 41 | 41 | 60 | 23 | 21 | 21 | 31 |
| 3 | 29 | 33 | 33 | 59 | 22 | 13 | 13 | 27 |
| 4 | 27 | 40 | 41 | 72 | 18 | 19 | 19 | 35 |

**Table 2.** Segmentation on non-standard data compared to wild type

|  | Jaccard | Q (C*A*L) | Connectivity | Area | Length |
| --- | --- | --- | --- | --- | --- |
| Wild type (n = 19) | 0.63 | 0.7 | 1 | 0.76 | 0.92 |
| Experimental conditions (n = 6) | 0.68 | 0.69 | 1 | 0.76 | 0.91 |

### Detection of angiogenic patterning changes with Vessel Metrics

Inhibition of VEGF through treatment with 10µM DMH4 from 24 to 75hpf (as compared to control 0.1% DMSO) is expected to reduce vessel growth. Six DMH4 treated and six vehicle DMSO treated embryos were imaged at 75hpf and analyzed using Vessel Metrics. Five parameters were compared between datasets. We find that branchpoint density and mean segment length were not significantly different suggesting that these fundamental properties of vessels are normal. However, we found statistically significantly reduced vessel density (p<0.05; two sided T test), network length (p<0.0001) and total segment count (p<0.0001) in VEGF-inhibited fish as predicted (Figure 4).

**Figure 4.**
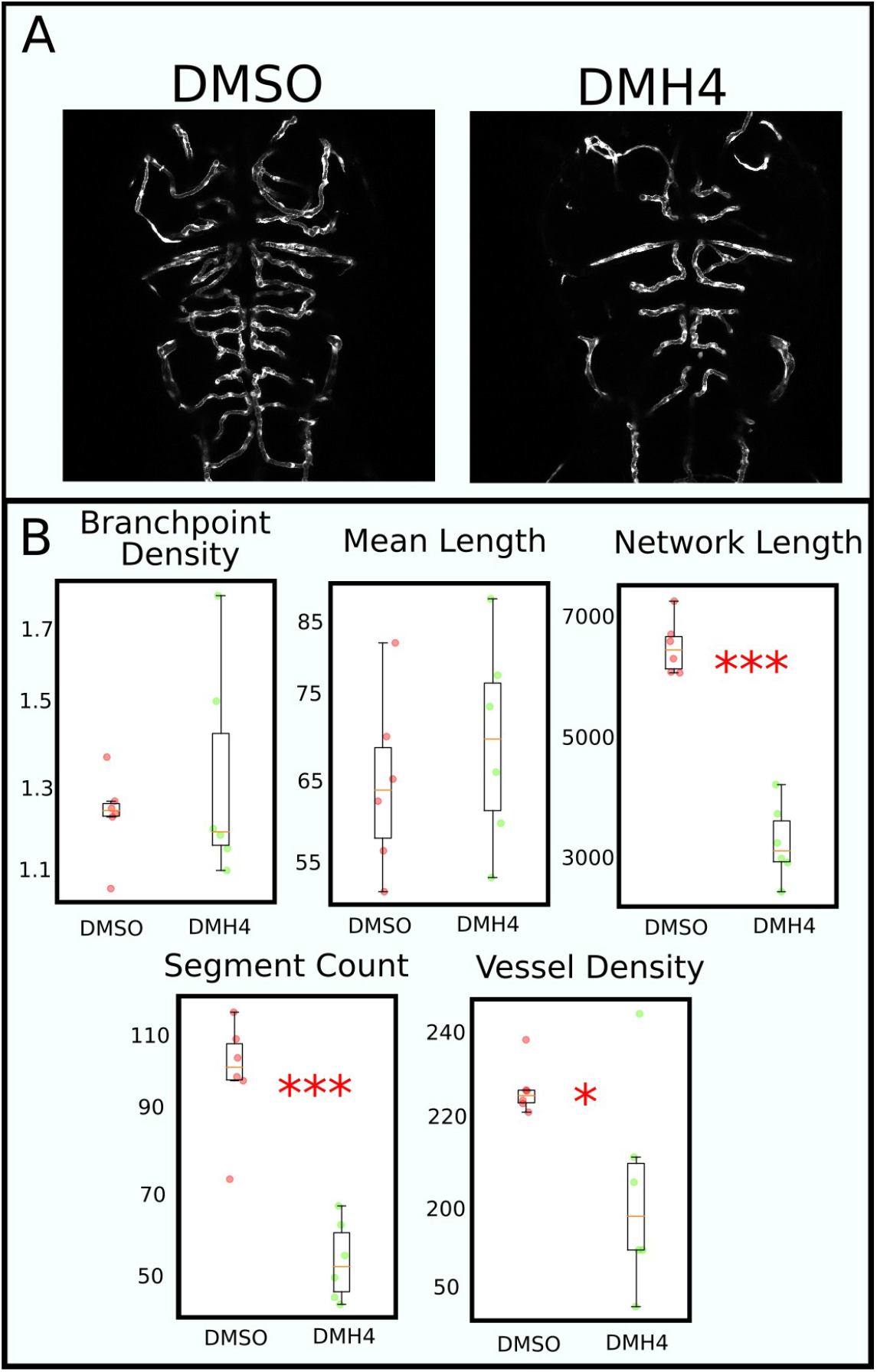
Parameter calculation of brain vessel properties after VEGF inhibition. A: sample image of DMH4 (n = 6) and DMSO (n = 6) (vehicle) treated fish. B: Scatter boxplots showing parameter distribution, one point per embryo. All p-values are calculated via 2 sided T test. Mean values (DMSO, DMH4) in pixels and p-values for each parameter: Branchpoint density (1.22, 1.31, p =0.48), Mean length (64.36, 69.36, p = 0.46), Network length (6480, 3640, p = 1.3E-6), Segment count (103.6, 54.5, p = 6.4E-5), Vessel density (227.1, 204.0, p = 0.037).

As a second proof of the sensitivity of the software, we tested whether the continuing development of embryonic brain vasculature, or changes in complexity could be detected by Vessel Metrics. Significant angiogenesis continues through development. We tested whether changes between images collected at 3dpf and 5dpf would reveal increased angiogenesis. Using five 5dpf and three 3dpf images, we calculated parameters using Vessel Metrics default settings. While branchpoint density, vessel density, and mean segment length were not different, we find that the network length (p<0.0001) and segment count were statistically significantly increased as would be expected for a developing embryo (p<0.05; 2 sided T test; Figure 5).

**Figure 5.**
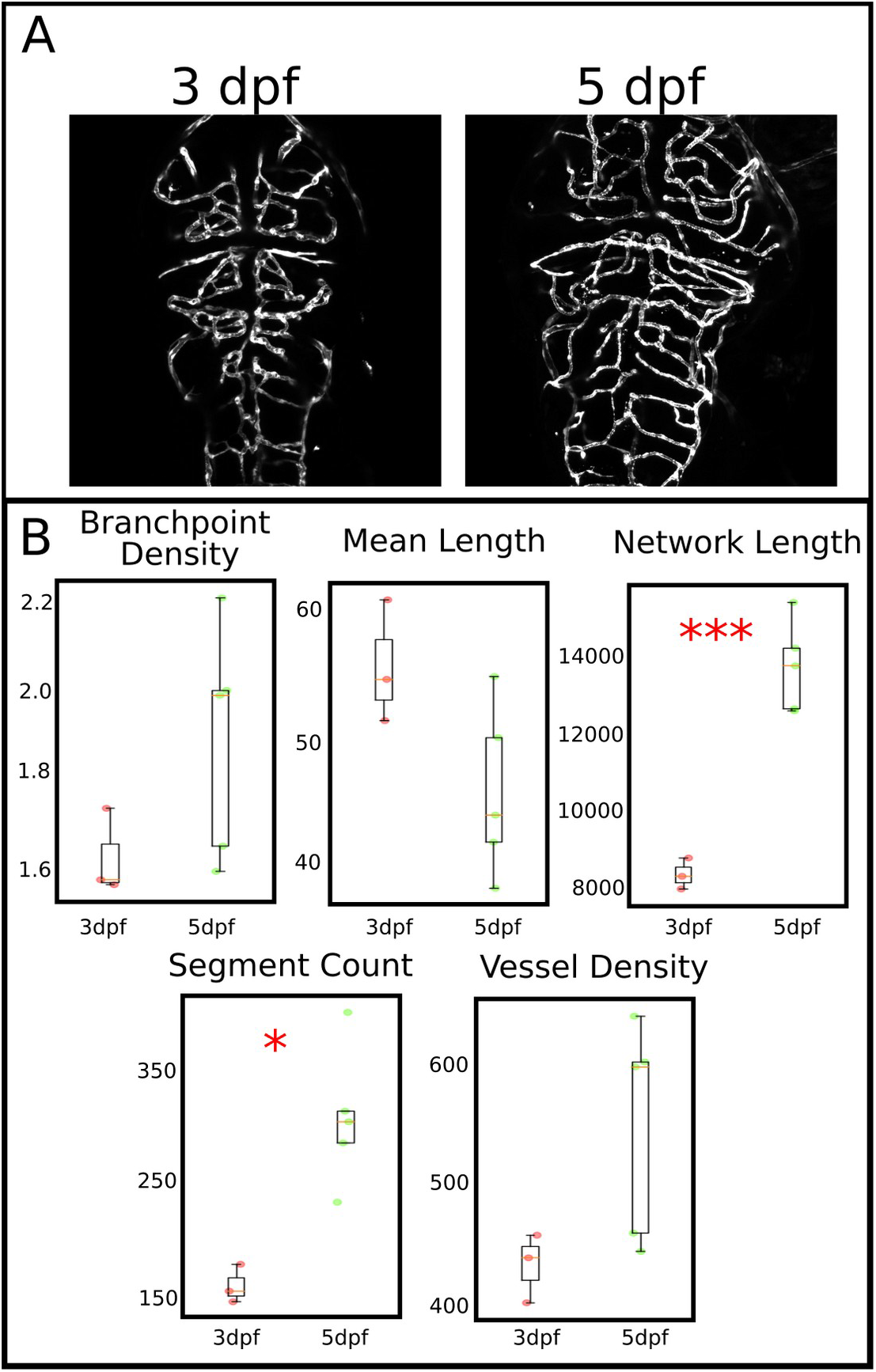
Parameter evaluation on 3 day vs 5 day zebrafish embryo. A: an example of the brain vasculature imaged dorsally of a 3 dpf (n = 5) and 5 dpf (n = 3) embryo. B: Quantification of Network length, segment count, and vessel density show significant increases in network length and segment count in 5dpf fish. All p-values are calculated via 2 sided T test. Mean values (3dpf, 5dpf) and p-values for each parameter: Branchpoint density (1.61, 1.87, p =0.15), Mean length (55.59, 45.53, p = 0.069), Network length (8400, 13730, p = 2.9E-4), Segment count (153.0, 309.4, p = 0.007), Vessel density (440.0, 554.1, p = 0.080)

### Vessel diameter comparison between manual and automated analysis

Vessel diameter is a critical measurement as it can differentiate vessels of different types (capillaries, arterioles, arteries etc.) as well as changes to vessels dynamically induced by external stimuli (increased heart rate, vascular relaxation or contraction). We took a unique approach to determining vessel diameter. We leveraged the image segmentation to determine individual vessel segments, then placed perpendicular crosslines along the vessel structure. Vessel Metrics removes the need to manually place cross lines or vessel centerlines. This method fully automates a previously developed semi-automated method (Figure 6). Additionally, a layer of error tolerance is built into Vessel Metrics with automatic outlier removal using a user specified tolerance. Removed measurements appear on the vessel visualization with a different color cross line so the user is aware of if or how many measurements were removed. In testing the software, we find that about 5% of crosslines were removed, primarily at the end of vessel segments where signal drops off or in locations where an adjacent vessel’s signal interferes. An example of diameter measurements with removed measurements shown can be seen in Supplemental Figure 3.

**Figure 6.**
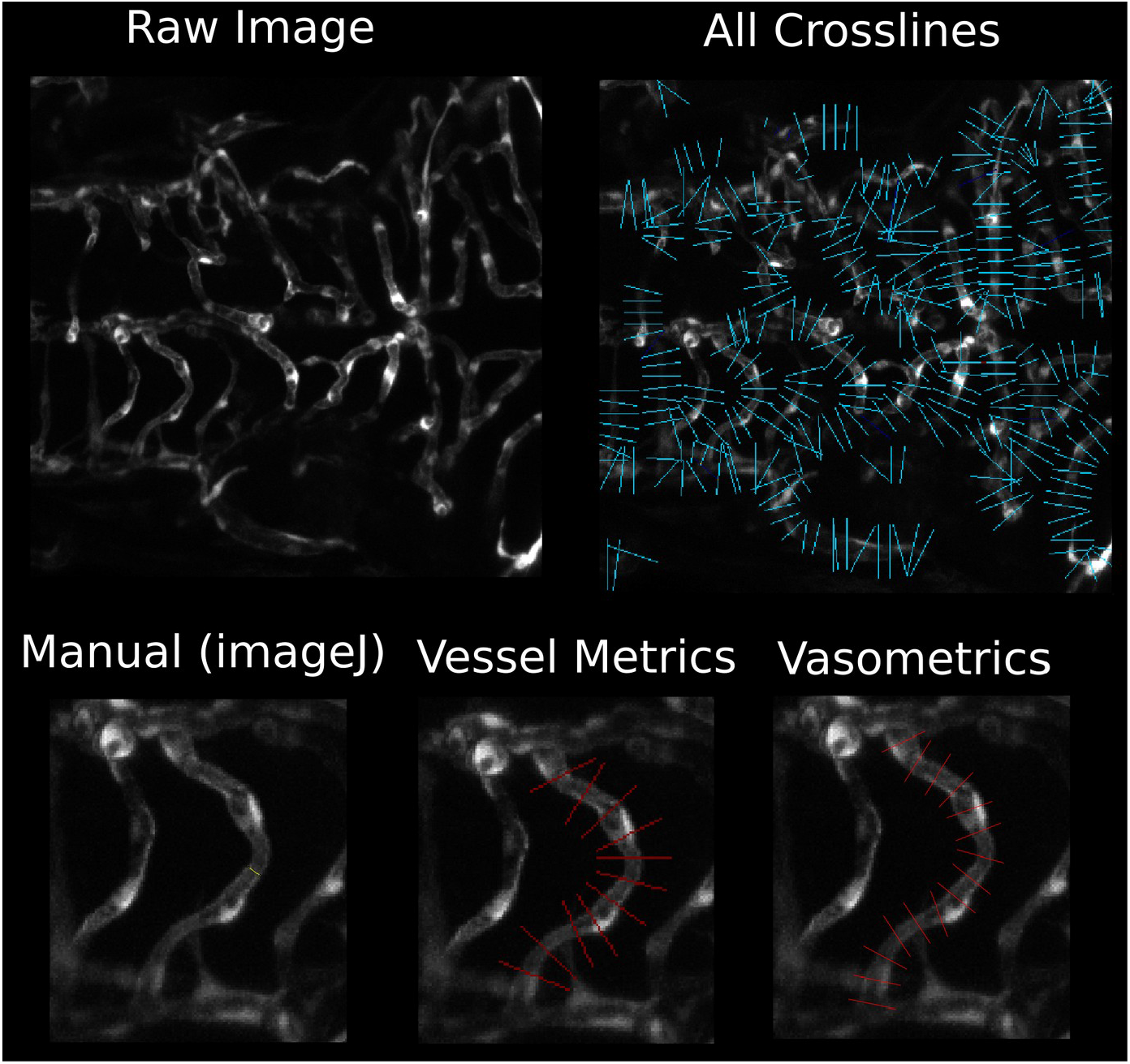
Comparison of methods to determine vessel diameter. Top left: raw image. Top right: all vessel diameters taken within a single sample. Measurements are generated in spreadsheet format with an average per segment and each individual measurement. Bottom: Demonstration of methods for vessel diameter comparison, manually using ImageJ, automatically with Vessel Metrics, and semi-automatically with Vasometrics.n = 25 segments, 5 segments from each of 5 embryo

Vessel diameter was compared from measurements made manually, semi-automatically using the Vasometrics app^17^, or automatically using a full width half max measurement on 25 segments (5 segments from each of 5 wild type embryo 3 dpf) in Vessel Metrics. For the manual measurement, three measurements were taken and averaged, one at the midpoint of the segment and one upstream and downstream. Automatically placing a larger number crosslines ensures vessel diameter changes along a segment are captured, and erroneous measurements (if undetected by the outlier detection) have a smaller impact on the average over the segment. We find that the mean percent difference between manual and Vasometrics was 27%. The mean percent difference between manual and Vessel Metrics was 10%. This data shows a substantial decrease in error, a vessel measuring 20 pixels across (common in the datasets used for this study) will have its error reduced from 5 pixels to within 2 pixels of the average of 3 manual measurements using Vessel Metrics.

### Application of Vessel Metrics to mouse brain confocal imaging

Brain blood capillaries in any species have a minimal diameter that is able to allow an erythrocyte to circulate, hence are relatively conserved across all vertebrates (∼5 μm on average). Imaging techniques vary across species, but confocal imaging is widely used in the mouse model. Confocal Z-stacks of murine brain vasculature (n = 5) were analyzed using Vessel Metrics. We find that only minimal adjustments are needed. We find that an additional step needs to be added to the segmentation algorithm in that a vessel enhancement filter at two scales needs to be applied to the images. These two enhanced images were combined (summed and normalized to 0-255) to achieve a suitable level of enhancement on both the microvessels and the larger vessels. The images were manually annotated and the accuracy of the segmentation algorithm quantified via Jaccard index and the vessel specific accuracy metrics. (Figure 7) The murine data had a mean Jaccard index of 0.67, and a Q (C*A*L) value of 0.7.

**Figure 7.**
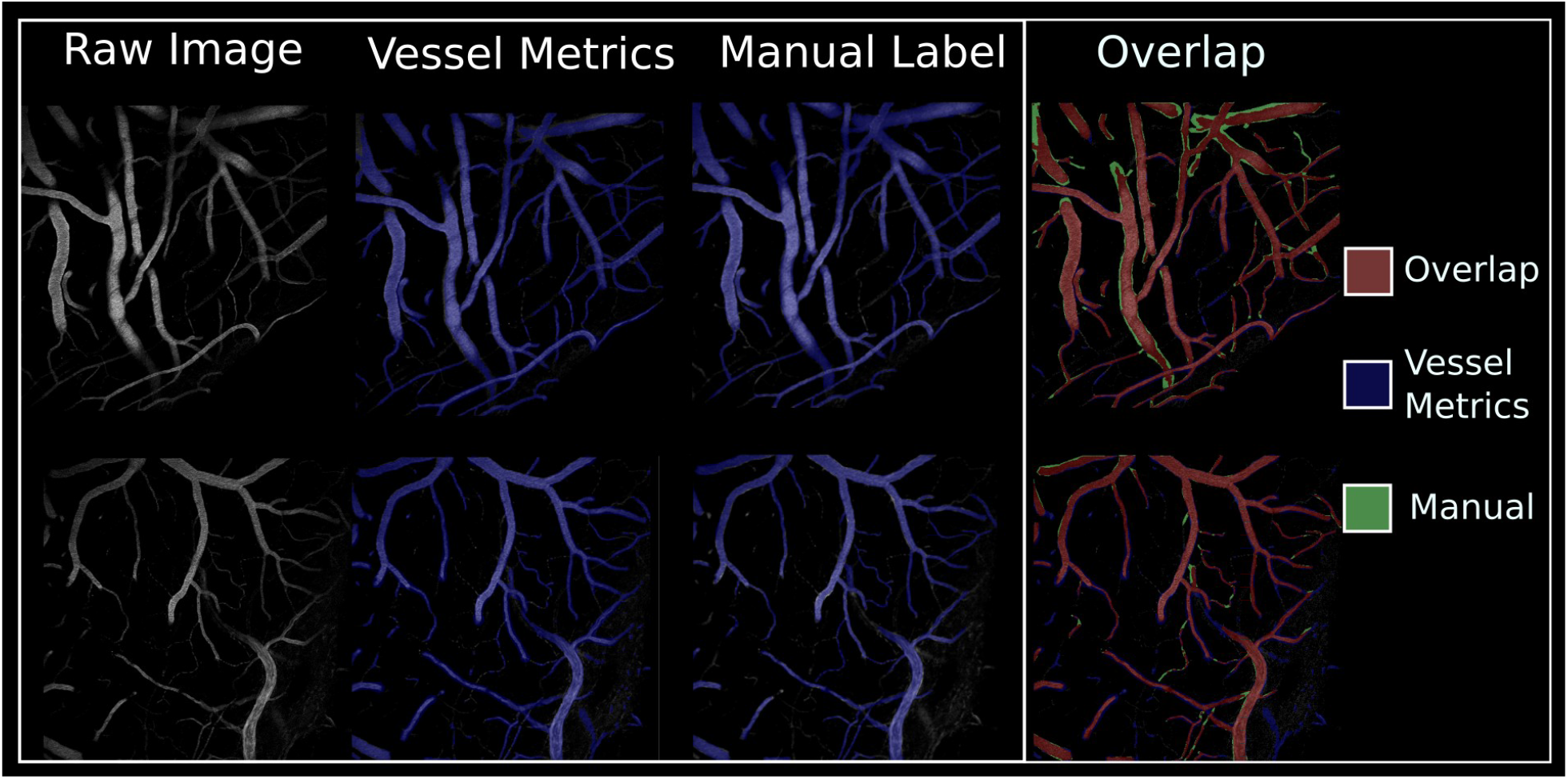
Vessel Metrics vs Manual segmentation comparison in mouse brain confocal images. (n = 5) Overlap image shown on the right, red pixels indicate agreement between manual annotation and Vessel Metrics.

Vessel diameter was also tested on the murine data. One segment from each image was selected, manually measured (mean of 3 measurements), and compared to the automatic vessel diameter output by Vessel Metrics. We found a mean percent difference between manual and automatic measurements of 4.8%. On vessels averaging 18.8 pixels in diameter this equates to less than a 1 pixel difference between manual and automatic measurements, suggesting that automated diameter measurements are highly accurate. This data also suggests that the technique is generalizable beyond zebrafish data.

### Comparison to Other Vessel Analysis Software

The REAVER dataset consisting of 36 light sheet microscopy images at 20x and 60x from six locations in a murine sample were used for this experiment. The segmentation outputs of AngioQuant, AngioTool, RAVE, REAVER, and Vessel Metrics were compared to manual annotations using a Jaccard index. REAVER out performed every other software tool on the dataset it was developed on. Vessel Metrics was the second most accurate software on this dataset. The average Jaccard index can be seen in Table 3.

**Table 3.** Segmentation accuracy of vessel analysis software compared to manual analysis on REAVER dataset.

| AngioTool | AngioQuant | RAVE | REAVER | Vessel Metrics |
| --- | --- | --- | --- | --- |
| 0.65 | 0.37 | 0.55 | 0.85 | 0.68 |

REAVER’s base segmentation algorithm was then applied to our zebrafish wildtype 1 (n = 10) and DMSO-treated fish from the WT/IWR dataset (n = 9). A Jaccard index was calculated and compared to Vessel Metrics. Vessel Metrics had an average Jaccard index of 0.67, REAVER had an average Jaccard index of 0.25. Representative segmentation results from REAVER and Vessel Metrics can be seen in Supplemental Figure 4. This result suggests that many programs will work the best for the data that they were developed on, however we have succeeded in making a vascular quantification software that is generalizable to different types of data from different species and gathered on different confocal microscope platforms, not only the zebrafish data that it was developed wtih. Only minimal user adjustment is needed when changing data types.

### User Interface and Output

A step by step, guided UI based method for setting up experiments is available for Vessel Metrics. A user can call the user interface script from the command line to be led through a series of steps to analyze a single image or directory of images. Users can save settings from previous experiments and load them later, so subsequent analyses require minimal time to run. Results are output in the user-specified directory with each image receiving a folder for its results, and images within the folder named identically. The processed images, labels and vessel centerlines, vessel labels, and density labels are saved along with a spreadsheet containing the user specified parameters. A user wanting to find the diameter of a specific vessel can find that information by finding the vessel segment label on the vessel label image and finding the diameter measurement corresponding to that segment number in the spreadsheet. Sample output images can be seen in Supplemental Figure 5.

## Discussion

Imaging data is rich in information that is often not captured in biological analysis. The development of simple, adaptable, accurate algorithms to extract information automatically has the potential to increase the ability to discriminate real differences in data that are not obvious by eye or hand measurement. In this manuscript, we detail the steps involved in the development and validation of the Vessel Metrics software for the analysis of vascular imaging. We outline the logic of the software development and the reasons for choices made in order to enable end users to customize the software to their own use. We demonstrate an accurate segmentation algorithm that identifies vessels from raw confocal images of transgenic zebrafish, transgenic mouse vessels, and light sheet mouse images. While our preprocessing steps are similar to prior, and contemporary work there are differences in vessel filters and downstream analysis that result in a highly consistent model across image modalities.^18,19^

Our goal in creating this software was to expedite the analysis of vascular images in the form they are currently analyzed, and provide consistency between research groups. Generating a maximum intensity projection is critical to our workflow; the low signal in any individual confocal slice, and resolution difference between planes make a 3D analysis with experimental confocal data infeasible. The 2D z-projection is imperfect but on the whole, represents the architecture quite well. Morphological changes will still be captured on images taken from the same perspective across multiple samples.^20^ As a result of the projection, structures that have a low expression of the fluorescent protein or exist only partially within the plane of the projection produce low signal vessels which are unmistakably on the image but without enough signal intensity to perform a meaningful analysis of structure.

Vessel enhancement filters are at the core of morphological approaches to segmenting vessel structures.^21^ Successful vessel enhancement should brighten vessel signal, including at branch points, while maintaining a reasonable estimate of vessel boundaries.^22^ Previous studies have primarily used the Sato filter.^23^ Of the four filters tested, we find that the Meijering filter performs the best on our data. The Frangi filter struggles at vessel junctions while the Sato filter has issues with nearby vessels.^24^. The Frangi filter produces a lower intensity enhancement and may be best suited for particularly bright vessels. The Jerman filter produces the most uniform contrast of all enhancement filters tested, but overestimates vessel boundaries which can cause downstream issues measuring vessel and branchpoint density. Empirically, we find the Jerman filter to be the most successful on images with low contrast, where a more dramatic enhancement allows lower intensity signals to be picked up. On low noise, high contrast images the Meijering filter is the most successful as the filter picks up dimly lit vessels and complex branch points without overestimating the boundaries of vessels with strong signals.

The selection of appropriate parameters will vary between datasets. We found the background suppression should be large enough to not not lose signal from large vessels but not so large that no signal is removed. In this case, a rolling ball filter with roughly ⅓ of the width of the image is ideal with a Gaussian blur roughly equivalent to half the diameter of the larger vessels in the image. The Gaussian smoothing should be as small as possible to homogenize the contrast within vessels as a large smoothing filter blurs the borders of vessels and leads to inaccurate diameter measurement and potentially erroneous branches in the vessel skeleton. An appropriate sigma range for the enhancement filter is likewise essential. The sigma range defines the diameters of tube shaped objects to enhance in the image (a range of 1-8 with a step size of 1 looks for vessels of diameters 1-8 pixels), choosing an inappropriate value will result in an inaccurate segmentation. Vessel Metrics was designed so users can adjust the parameters of the analysis pipeline to their own workflow for each individual dataset. The major parameters affecting segmentation output are tunable via the user interface. The interface suggests default values which perform well on all data included in this manuscript.

As an added test of functionality, we extended analysis of Vessel Metrics functionality on murine data, to provide evidence in support of generalizability. The murine data displays inherently different image characteristics compared to zebrafish data: there is a much larger variance in vessel size and the image is noisy with a smaller variance in pixel values. The preprocessing method developed on zebrafish smooths the noise and amplifies vessel signal, allowing for an effective segmentation. The murine data included a wider range of vessel thicknesses which resulted in overestimation of vessel boundaries on small vessels. A multi-scale segmentation technique used for the murine data was implemented to combat this problem. We found this step was not necessary for the zebrafish images because the vessel sizes are more uniform. In zebrafish, confocal images vessels range from 2-15 pixels in diameter, whereas in the murine data vessels range from 2-50 pixels in diameter. This greater variance requires additional steps to successfully characterize all sizes of vessels. The multi-scale segmentation was likewise used with default parameters for the REAVER dataset comparison.

The complex and irregular nature of vasculature imaged with confocal microscopy lends itself to imperfect skeletonization. Correctly identifying individual segments and branch points is critical to downstream parameter analysis. Vessel Metrics introduces post processing steps unique to vascular analysis to clean up the raw skeletonization and produce a more accurate structure. Small terminal segments typically caused by active angiogenesis and irregularities in vessel shape are removed and long segments are combined. Erroneous branch points are deleted and the connected segments are corrected. We found that implementing this post processing reduced our average percent error compared to manual measurements by over 50%. Additionally, the region cropping feature can be used to remove sections of the image with aberrant signals or focus on a particular area of interest.

We tested inter-user variability as a means to validate the reliability of our segmentation technique. Three observers annotated wild type data and the Jaccard index was compared between pairs of human observers as well as human observers and the algorithm. We found two primary regions of difference between observers: in annotating abnormal vessels and vessel boundaries. Abnormal in this context broadly describes vessels with strange contrast or boundaries which are (to a human eye) clearly not useful for analysis. Some observers chose to annotate these vessels regardless and others chose not to. Similarly, vessels with low signal due to partial voluming or poor expression of the fluorescing protein were excluded by some observers. Observers tended to disagree regarding vessel boundaries. Due to the shape of the vessels, a disagreement of 1-2 pixels along the vessel boundaries for major vessel segments can skew overlap based metrics substantially. We found the mean agreement between observers to be nearly identical to the mean agreement between any observer and Vessel Metrics, indicating the algorithm is as good as a human observer with the added benefit of computational speed and reproducibility.

Determining accurate microvessel diameter is one of the most useful parameters in Vessel Metrics, where experimental vessel diameter changes may differ by as little as a pixel or two (for instance during experiments inducing constriction and relaxation of vessels). A comparable existing ImageJ plugin, Vasometrics, requires the user to draw a vessel centerline before the software places evenly spaced cross lines perpendicular to the user drawn line and measures diameter using a full width half max measurement. Vessel Metrics has improved the accuracy of cross line length determination by leveraging the vessel segmentation. The vessel label is a more stable means to determine the size of a vessel segment whereas placing a test crossline on the raw image may provide an inconclusive measurement. Error tolerance is built into Vessel Metrics; after diameter at each crossline is measured, measurements outside a specified tolerance are discarded and the crosslines marked to indicate to the user which measurements likely contained errors. Vasometrics requires the user to manually identify and remove outlier measurements and relies on raw image signal to suggest cross line length, which fails about 50% of the time in practice. Vessel Metrics similarly discards vessel segments with a poor segmentation, where an automatic crossline determination isn’t feasible. Vessel Metrics requires less user input, provides data on the entire vessel structure, and produces results which are closer to manual measurements.

Other vessel detection software is available and unfortunately underutilized. Commercial software is available with select microscopes such as Nikon’s NIS-elements (www.nis-elements.com); but is not open source. Kugler et al. produced a segmentation technique specific to zebrafish vasculature in light sheet microscopy.^4^ Additionally, a deep learning technique to segment vasculature on confocal imaging has also been published.^25^ In terms of end to end vessel structure analysis, semi automated analysis is possible through various imageJ plugins such as Vasometrics^17^, or using open sourced packages such as AngioQuant^26^, RAVE^11^, AngioTool^27^, Zebrafish Vascular Quantification (ZVQ)^19^ and REAVER.^18^ REAVER and ZVQ are the most recent comprehensive vessel analysis tools. REAVER uses a more fine tuned segmentation method than the other tools, and is capable of outputting a number of useful parameters. However, despite the availability of tools, a recent report noted an order of magnitude difference between the number of manuscripts quantifying vessel structure and citations of this vessel detection software.^3^ Clearly there is a problem in widespread usage across the field, even though standardization of reporting of vessel parameters is ideal. We would anticipate that there is a large inter-user inter-lab variability with respect to vessel parameters. We attempted to quantify the inter- and intra-user variability across a variety of conditions and microscope parameters and found that the software performed equally, suggesting that automation of this process could eliminate much of the variability in the field. Quantifying the effect size and thus the need for software tools is a critical step in standardizing the methods used to analyze vascular images.

We found a favorable comparison to the segmentation algorithms available through other freely available software tools. Vessel Metrics segmentation algorithm performed second best on REAVER’s benchmark dataset. Further, across all datasets tested Vessel Metrics mean Jaccard index fluctuates between 0.65-0.7. Vessel Metrics performs consistently across datasets and animal models, which is a highly valuable trait for a general use software tool. Although machine learning is being used more frequently, on a dataset by dataset basis fully training a deep learning model will provide higher base accuracy at the cost of requiring a larger, curated training dataset. For biologists quickly taking measurements on a small sample of complex images a consistent generalizable tool is more likely to be used than a higher accuracy tool that requires more input time to validate and test it on their particular data type.

Deep learning has been successfully used to segment similar images in the past^25^, and on a dataset by dataset basis the accuracy metrics are higher than what’s achievable with a traditional image processing algorithm. However deep learning models tend to be dataset-specific and generalization to a broader set of data may require additional processing and training.^28^ In terms of creating a generalizable set of tools to segment and process a variety of image types, a traditional image processing algorithm such as Vessel Metrics still plays an important role. A deep learning model is only as good as the dataset it was trained on, and will have variable success on images of the same type acquired at different resolutions or on a different microscope, and likely need retraining with each additional type of data. Depending on the dataset and the skillset of the scientists asking the research question this may be prohibitively intensive and not feasible. A set of image processing algorithms which find the major vessels and automatically analyze the data may have uses still despite a lower accuracy simply due to ease of adoption and customizability.

In conclusion, we present Vessel Metrics, a software tool for the automated analysis of vascular images with a focus on confocal microscopy, that is an effective tool for comparing different developmental stages as well as disease or mutant states for accurate measurement of phenotypes. Vessel Metrics allows for fully automated, tunable analysis of large microscopy datasets in a manner that is reproducible and generalizable. Our software fully automates diameter measurement with built in error tolerance. Vessel Metrics outputs a standard set of parameters that can distinguish vascular growth and disease. We’ve demonstrated that the segmentation matches human observer quality and is functional on data collected on multiple microscopes, at varying stages of development, and on two species. It is our hope that the adoption of Vessel Metrics reduces the time to analyze experimental results, improves repeatability within and between institutions, and expands the percentage of a given vascular network analyzable in any given experiment.

## Methods

### Code and Data Availability

Vessel Metrics source code is available at: https://github.com/mcgarrysd/vessel_metrics. Vessel Metrics was written in python 3.8 and requires standard python libraries as well as the EasyGui, bresenham, and aiscimageio libraries. The image datasets and annotations used in this manuscript are available via github.

### Animal Care

All experimental procedures were approved by the University of Calgary’s Animal Care Committee (protocol AC21-0169). Transgenic *Tg(kdrl:mCherry)*^*ci5*^ and *Tg(kdrl:EGFP)*^*la116*^ zebrafish embryos were maintained at 28.5C in E3 medium until imaging at 75hours post fertilization.^29,30^ Male transgenic Aldh1l1-Cre/ERT2 x RCL-GCaMP6s P22-60 c57bl/6 mice were imaged for the murine dataset (data kindly provided by Dr. Grant Gordon, protocols AC15-0133 and AC15-0053).

### Image acquisition

Zebrafish embryos were raised at 28C to 75 hours post fertilization and imaged using a 20x objective (NA = 0.8) on a Zeiss LSM 880 or LSM 900 Confocal Microscope while mounted in low-melt agarose. The murine data was imaged on a custom two -photon microscope with a 40x objective, NA = 1.0 processed in ScanImage running in MATLAB.^31^

### Datasets

Four datasets were used in this study. Wildtype 1 consists of 10 wild type embryo dorsally imaged zebrafish whole embryonic brain images cropped at the midbrain. WT/IWR consists of 9 wild type embryos and 8 embryos treated with the small molecule inhibitor 10µM IWR1 (Sigma Aldrich, Cat I0161). DMSO/DMH4 includes 12 embryos at 3 dpf, 6 treated with 0.1% DMSO (Sigma Aldrich, Cat D8418) and 6 treated with 10µM DMH4 (Sigma Aldrich, Cat D8696). Wildtype 2 includes 5 wild type embryos at 5 dpf and 5 wild type embryos at 3 dpf. The murine dataset includes 5 wild images, the murine dataset is a full z projection rather than the half z projection used for the other datasets. A summary of datasets is found in Table 3.

**Table 3.**
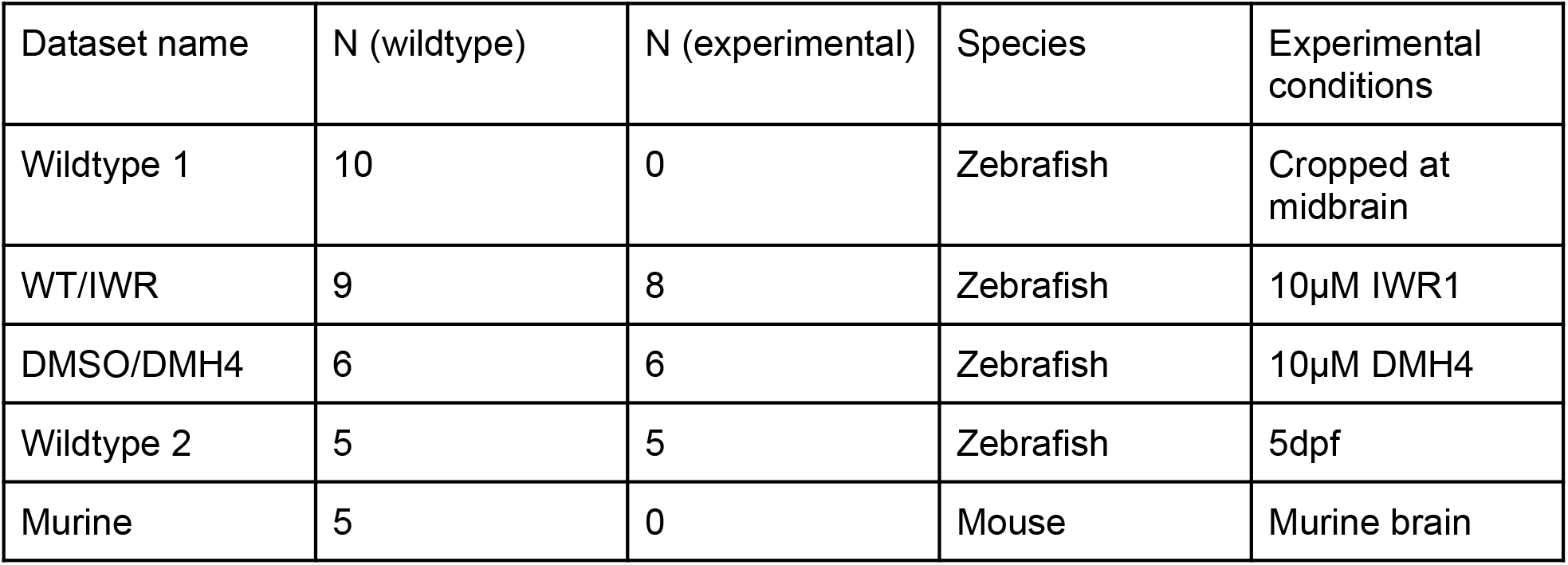
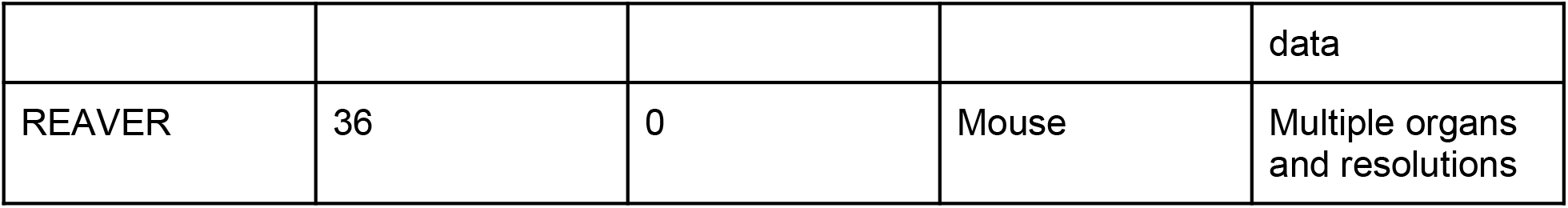
Summary of datasets

### Image preprocessing and segmentation

Images were loaded into python in native czi format using the czi file library. The channel corresponding to the labeled endothelial cells is selected and the image is normalized to an integer range from 0-255. A maximum intensity projection is calculated over half of the image stack. For this manuscript the bottom half of the stack is used. A linear contrast stretch is applied to the image using the 5th and 95th percentile values using equation 1 below.^32^

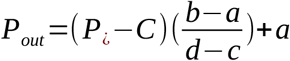

In this equation, Pout is the new pixel value, Pin is the input pixel value, a and b are the lower and upper limit pixel values (0 and 255 in the case of this manuscript) and c and d are the 95th and 5th percentile pixel values respectively.

A gaussian blur (user defined size, default = 7) is applied to smooth inhomogeneities within the vessel, followed by a rolling ball background suppression filter with a size equivalent to ⅓ of the image.^33^ A user-selected vessel enhancement filter (Sato, Frangi, Meijering, Jerman) is applied with a user-selected sigma range (default Meijering filter with sigmas 1:8).^22,34–36^ The enhanced image is normalized and thresholded (default = 40). A morphological opening is performed on the label, small holes are closed, and large background artifacts removed to produce the final segmentation label.

### Wild type segmentation

The segmentation algorithm was tested on the wild type images from datasets Wildtype 1 and Wildtype 2 (n = 19). Ground truth values were created by SM using LabelMe (https://github.com/wkentaro/labelme). Overlap was calculated using a Jaccard index, as well as three vessel specific accuracy metrics, connectivity, area, and length.^13^ All four vessel enhancement filters were tested with varying thresholds to characterize the differences between the enhancement filters.

Connectivity, area, and length were calculated using equations 2-4 below respectively:

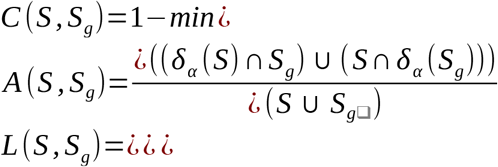

Each parameter is bounded between 0 and 1, where 1 is a perfect segmentation. S denotes the segmentation and Sg denotes the ground truth segmentation. #c denotes the number of connected components. δ denotes a dilation of the segmentation to control for variance in vessel boundaries. *φ* denotes a skeletonization and *δ* _*β*_ denotes a dilation of the skeleton to control for variance in vessel centerline between segmentations.

The three metrics can be combined into a single quality metric by multiplying the three together. The quality metric Q ranges from 0-1 with 1 being ideal.

### Segmentation versatility

The segmentation algorithm was tested on a small sample of drug treated fish (IWR and DMH4) and on 5dpf data (n = 2 each condition, n = 6 total). Data was manually labeled by SM using LabelMe.

Additionally, a sample of wild type 3dpf data (n = 5) was annotated by an additional 2 lab members (CA, IV).

### Parameter derivation

Vessel density was calculated by splitting the image into 256 evenly sized tiles and summing the number of vessel pixels by the total pixels within each tile. Vessel density is user customizable with respect to the number of x and y tiles used in each image. Users can choose to omit tiles with no vessels from the mean vessel density calculation.

The vessel label was skeletonized using the skimage library. Branch points were identified via a neighborhood analysis. For each pixel on the skeleton the sum of the immediate 3×3 neighborhood is calculated. Straight vessel segments have a neighborhood sum of 3, whereas branch points have a sum >3. The branch points were subtracted from the skeleton and each independent segment assigned a unique number. Segment endpoints were identified using an identical method to the branch points.

Each branch point breaks the skeleton into distinct segments, as a result large vessel segments with multiple child branches are broken into separate segments. In order to correct this, a neighborhood analysis was performed, identifying the segments which connect to a branch point. The slopes of these segments were measured from endpoint to endpoint. Segments connected to the same branch point with similar slopes were connected and the branch point amended, such that parameters could be taken over the entire length of the large segment.

Effective segment length was calculated by summing the number of pixels within the skeleton of each segment. Segment tortuosity was calculated as the chebyshev distance between end points over the effective segment length. Effective segment length and tortuosity can be output as Total network length was calculated by summing the number of pixels in the vessel skeleton.

### Parameter validation

In order to validate parameter accuracy, central sections were cropped from 4 DMSO treated fish from the DMH4/DMSO. Number of segments of branch points were counted manually. The Vessel Metrics skeletonization and parameter calculation were applied to both the manual label as well as the automatically generated segmentation.

In order to demonstrate the utility of automatic parameter calculation, vessel parameters were calculated on the DMH4/DMSO and 5dpf datasets. Parameters were calculated as a mean parameter per image. Mean parameter differences were quantified via 2 sided t-test, comparing drug treated to control fish.

### Vessel diameter calculation

Vessel diameter calculation is calculated segment by segment basis. For each segment, the mid point is first identified by calculating and sorting the euclidean distance between each pixel and a single end point of the segment; the midpoint is defined as the pixel with the median euclidean distance. The slope of a tangent to the segment midpoint is calculated using a 5×5 neighborhood with the segment midpoint as the middle pixel. The slope is determined by the endpoints within the neighborhood. A perpendicular slope is calculated by inverting the tangent slope. A preliminary cross line is placed through the segment midpoint with the calculated perpendicular slope. Appropriate crossline length is determined by applying a 2.5x multiplier to the width of the vessel label along the preliminary crossline. This multiplier serves to account for vessel diameter changes along the length of the segment. Cross line length is static for each vessel segment.

A similar procedure is used to automatically calculate vessel diameter along the segment. At 10 pixel intervals along the segment the tangent and perpendicular slope is identified, a cross line of predetermined length is placed, and the pixel intensity of the pre-processed image (denoised, smoothed, contrast stretched, without a vessel enhancement filter) is measured along the cross line. A full width half max distance is calculated as the diameter of the vessel at that point.

Erroneous data points are identified and excluded prior to outputting a mean diameter measurement for the segment. Segments with fewer than 5 measurements are dropped from the dataset.

Vessel diameters are output in table format along with an image displaying which vessel segments correspond to each segment number within the table. Additionally, a visualization of the cross lines across each segment can be generated.

### Vessel diameter validation

Vessel diameter was measured manually on 5 segments from 5 embryo (n = 25) in Fiji using the line drawing tool. Manual measurements were collected at 3 locations along the vessel segment: the vessel center, and one region on either side of the vessel center. Measurements were likewise taken on the same segments using the VasoMetrics Fiji plugin as a comparison to a pre-existing semi-automatic method.^17^ VasoMetrics measurements were calculated at 10 pixel intervals using the automatic crossline length detection. A cross line length representative of the vessel size was manually input if the automatic detection failed. Vessel diameter was measured both using the vessel label and using a full width half max measurement on the pre-processed image.

### Murine Data Analysis

Deep Confocal Z-stack images of murine brains were provided by GG. Five samples were used to validate the efficacy of Vessel Metrics on an alternative data type. Samples were annotated manually to create a ground truth comparison. Images were pre processed identically to the zebrafish data. A multi-scale segmentation approach was used to account for the more varied vessel sizes in the murine data; after the image was preprocessed two vessel enhancement filters are applied with different sigma ranges. One sigma range covers microvessels and the second sigma range covers larger vessels. These enhanced images were summed, normalized, and thresholded identically to the zebrafish data. The segmented results were compared to the manually annotated images via Jaccard index.

The automatic diameter code was similarly tested on the murine data. Five segments (one segment per image) were manually measured using ImageJ. The mean of three measurements across the segment was used as the manual diameter measurement.

### Reaver Comparison

REAVER’s benchmark dataset (n = 36) was used for this comparison. This dataset contains murine vessels from the diaphragm, heart, inguinal fat, liver, peritoneum, and retina at 20x and 60x fields of view. The images were acquired using lightsheet microscopy. The authors manual annotations and the segmentation outputs of RAVE, AngioQuant, AngioTool, and REAVER were included in this dataset.

Vessel Metrics was applied to the base images using the default segmentation algorithm parameters (the multi-sigma segmentation was used for this analysis to account for the varying vessel diameters in the image). The Jaccard index pairing each vessel analysis software with the manual ground truth was calculated.

REAVER’s segmentation algorithm was applied to the benchmark dataset from Vessel Metrics (n = 19, 75hpf DMSO treated). The Jaccard index pairing REAVER with manual annotation was compared with Vessel Metrics.

## Acknowledgements

Operating funding was provided to SJC by NSERC Discovery Grants RGPIN/07176-2019. Trainee funding was provided by the Libin Cardiovascular Institute Masters (CA) and Postdoctoral Fellowships (SM), Eyes High International Doctoral Scholarship (CA) and Postdoctoral Fellowship (SM) from the University of Calgary, Cumming School of Medicine Postdoctoral Fellowship (SA, SM), and Alberta Children’s Hospital Research Institute Postdoctoral Fellowship (SA), and Cumming School of Medicine Graduate Scholarship, Canadian Institutes of Health Research Canada Graduate Scholarship, University of Calgary Faculty of Graduate Studies Doctoral Scholarship, and Alberta Graduate Excellence Scholarship (JGW). We thank Dr. Grant Gordon and lab members Adam Institoris and Milene Vandal for the generous provision of their raw mouse imaging data.

## Supplemental Information

**Supplemental Table 1.**
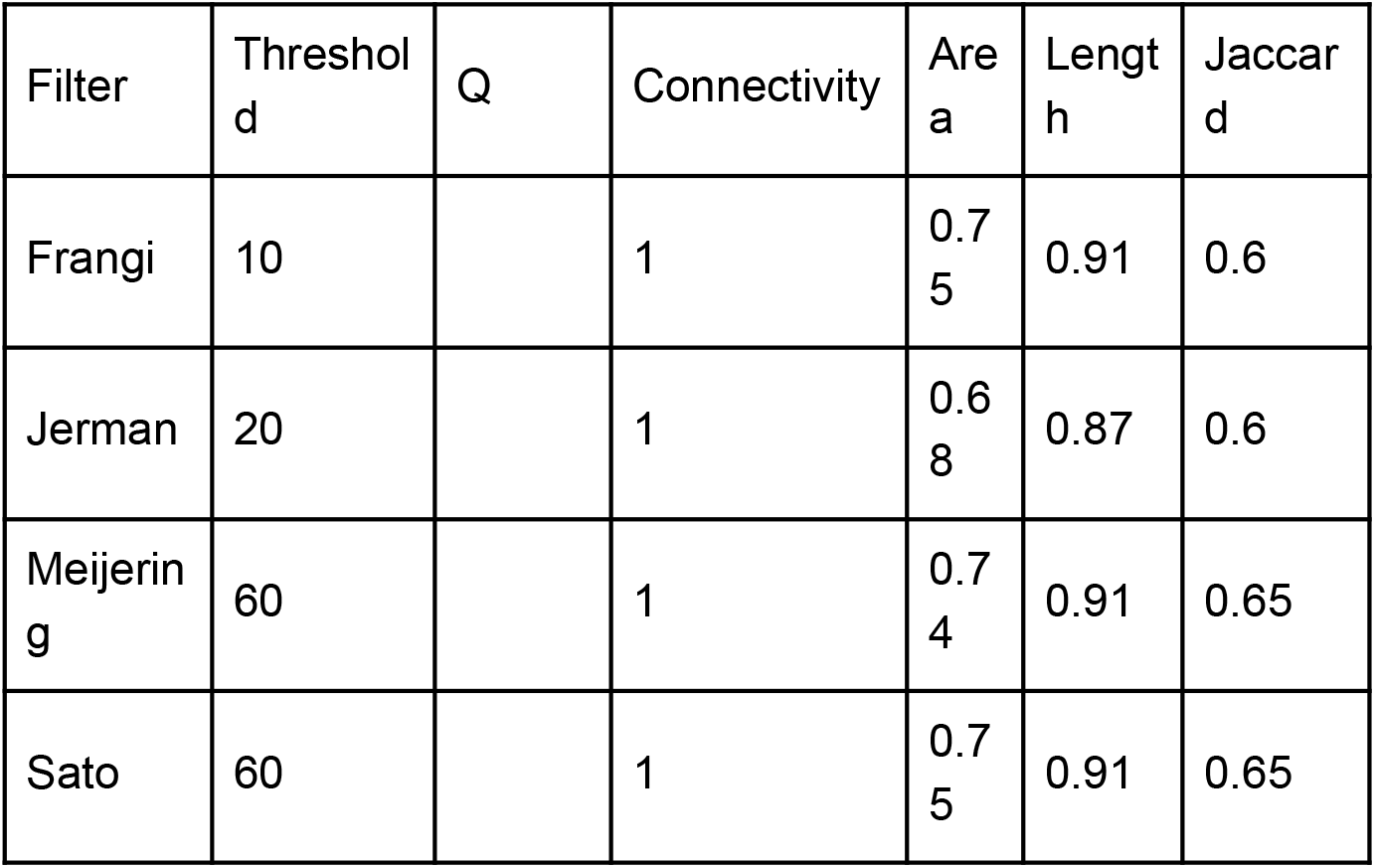
Segmentation results using other enhancement filters.

**Supplemental Figure 1.**
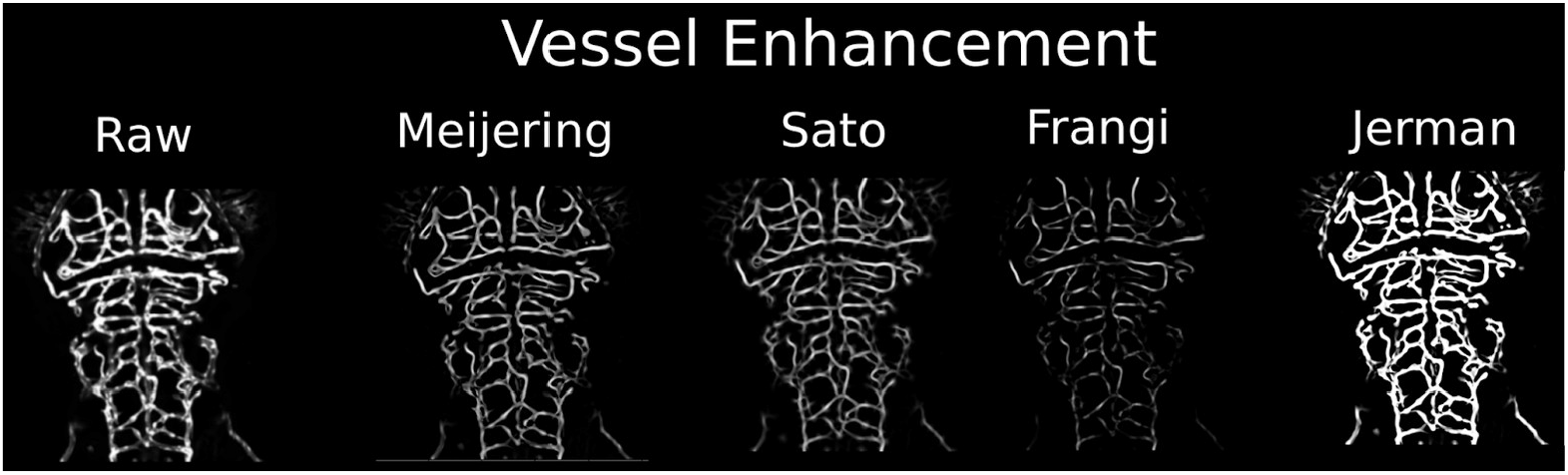
Examples of the four vessel enhancement filters applied to the same image.

**Supplemental Figure 2.**
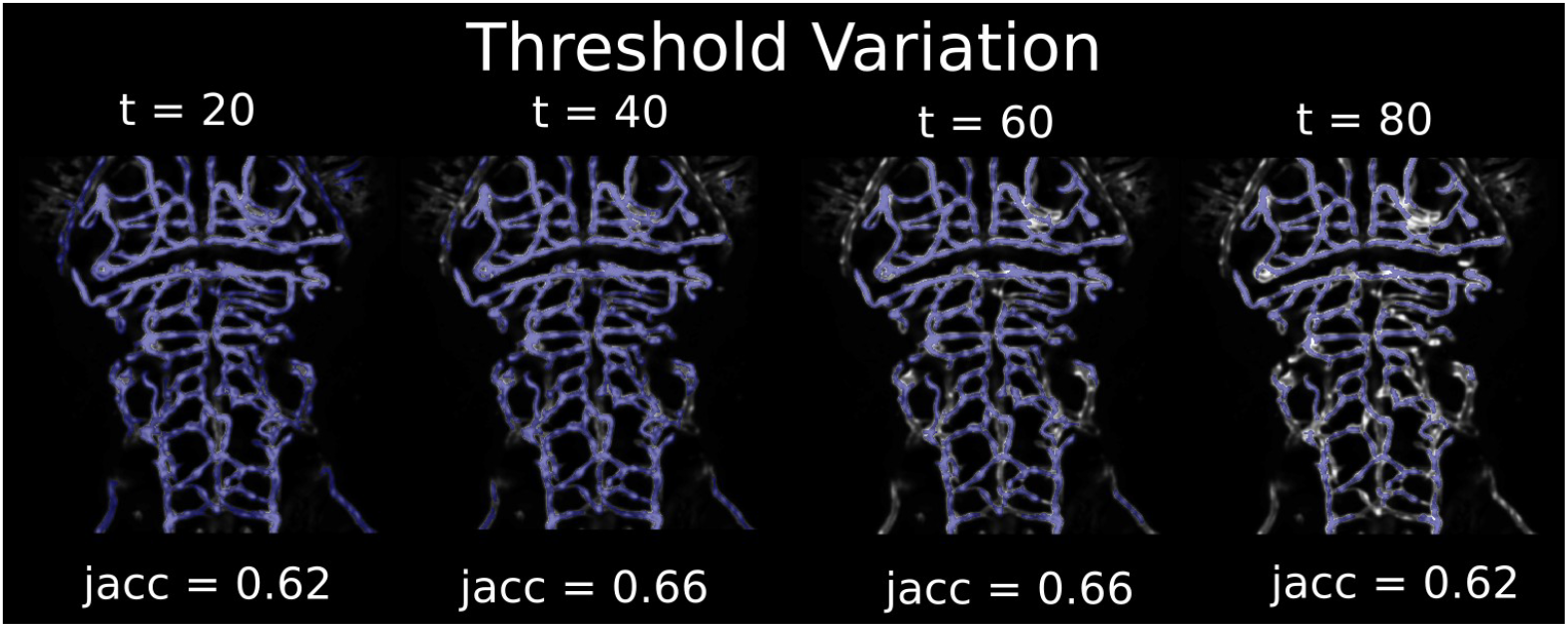
Effect on Jaccard Index by varying the threshold on the same image enhanced to the indicated values with the Meijering filter.

**Supplemental Figure 3.**
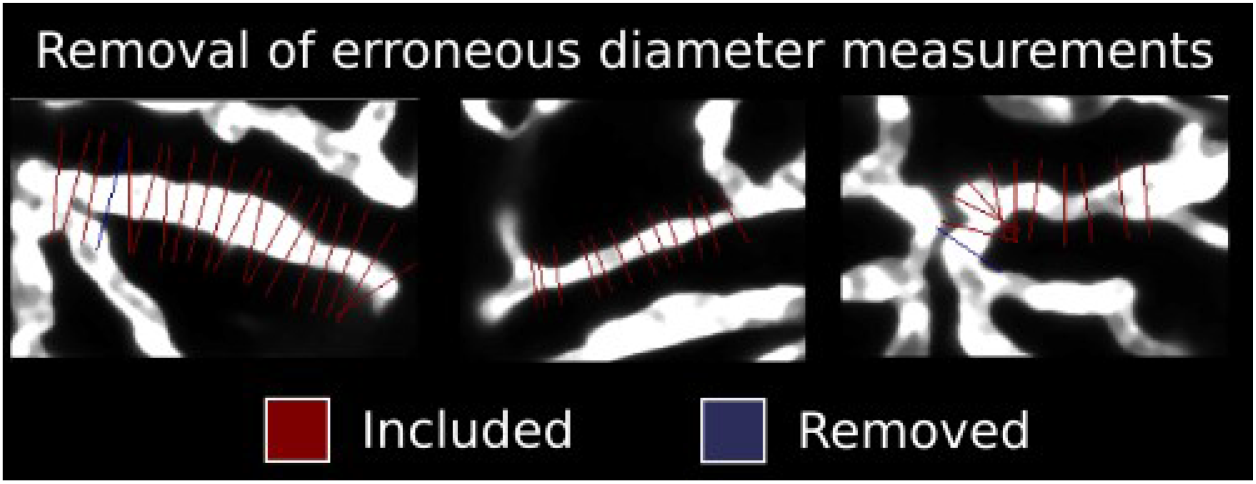
Sample diameter measurements. Measurements are removed when the image profile along the crossline is bimodal or resembles a step function. Removed measurements are shown in blue.

**Supplemental Figure 4.**
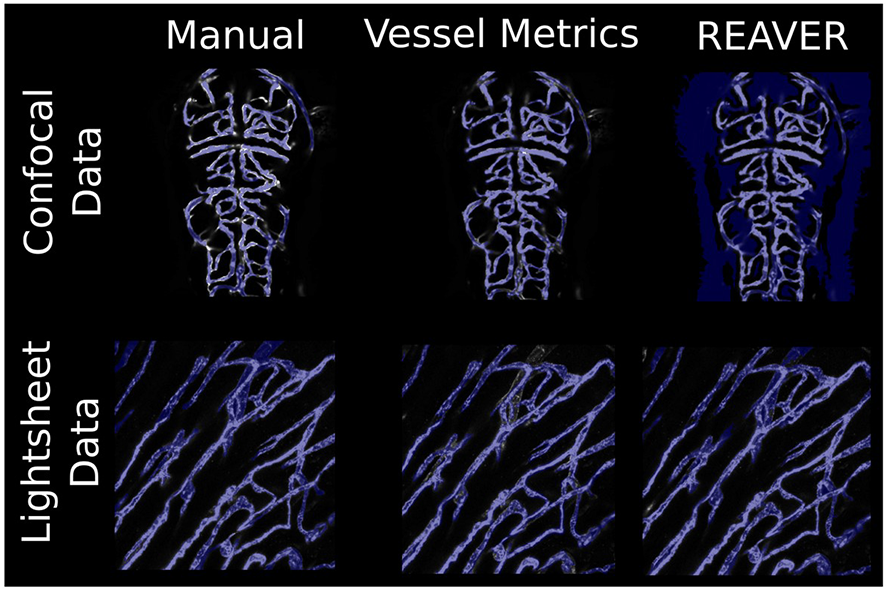
Segmentation comparison between REAVER and Vessel Metrics on confocal and lightsheet data.

**Supplemental Figure 5.**
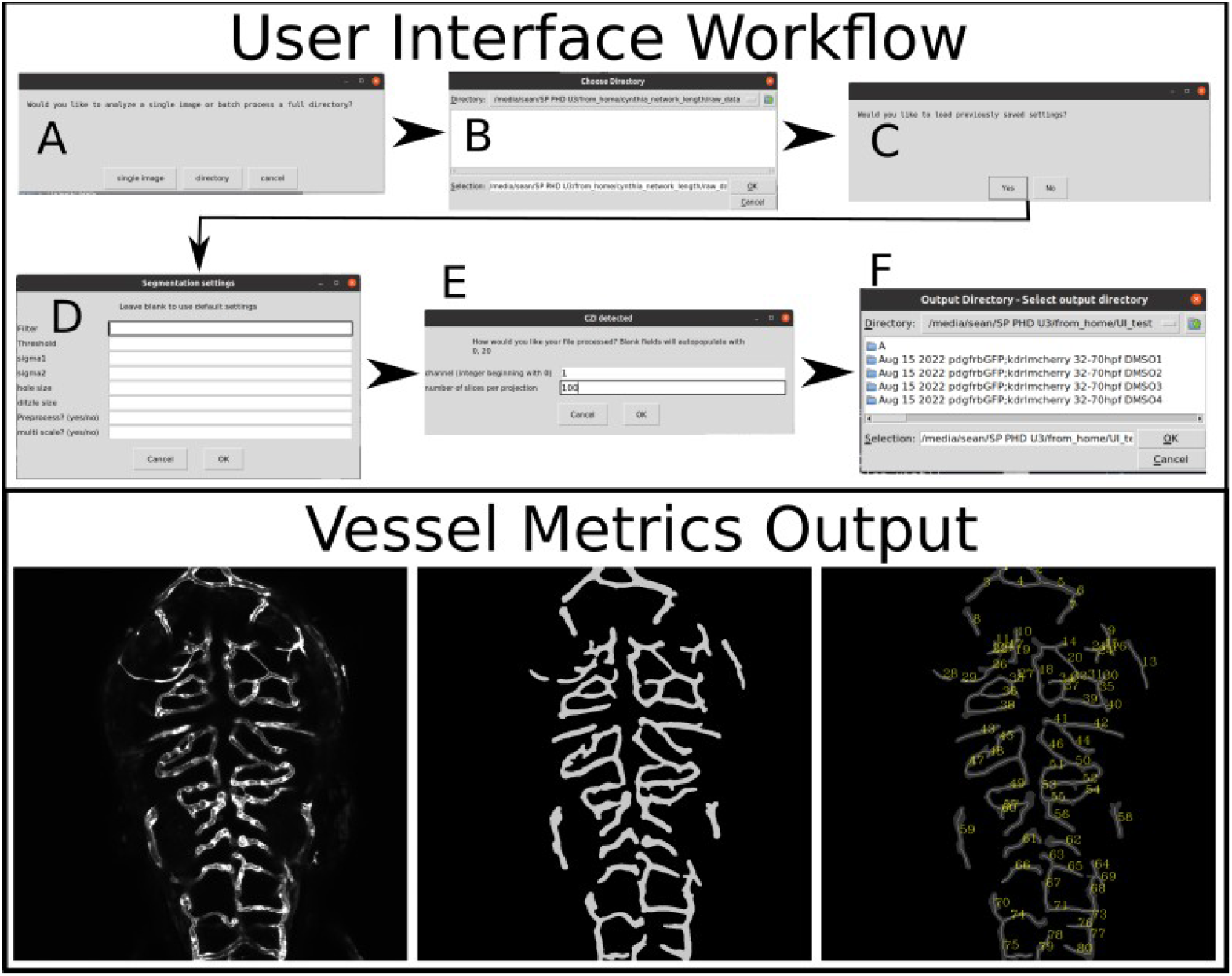
Top: User interface workflow. A. User can analyze a single image file or batch process a directory. B. Directory selection menu. C. Previously saved settings can be loaded to allow easier implementation of the same settings between experiments. D. Parameter values, fields left blank autocomplete with default values. Explanation of each parameter can be found in the readme. E. If a czi file is detected options to preprocess are presented, if the file is a single image this step is skipped. F. Output directory is selected. Bottom: Output of vessel metrics. Left: Z projection with no other processing. Middle: Segmentation right: Segmentation and vessel centerlines with vessel segment numbers. These labels match the parameter outputs in the accompanying spreadsheet.

## Notes

**Conflicts of Interest:** All authors declare no conflicts of interest in the publication of this manuscript

### Competing Interest Statement

The authors have declared no competing interest.

